# Gas6 ameliorates intestinal mucosal immunosenescence to prevent the translocation of a gut pathobiont, *Klebsiella pneumoniae*, to the liver

**DOI:** 10.1101/2023.01.19.524842

**Authors:** Hitoshi Tsugawa, Takuto Ohki, Shogo Tsubaki, Rika Tanaka, Juntaro Matsuzaki, Hidekazu Suzuki, Katsuto Hozumi

**Affiliations:** Transkingdom Signaling Research Unit, Division of Host Defense Mechanism, Tokai University School of Medicine, Isehara, Japan; Department of Hand Surgery, Nagoya University Graduate School of Medicine, Nagoya, Japan; Department of Immunology, Division of Host Defense Mechanism, Tokai University School of Medicine, Isehara, Japan; Division of Pharmacotherapeutics, Keio University Faculty of Pharmacy, Tokyo, Japan; Division of Gastroenterology and Hepatology, Department of Internal Medicine, Tokai University School of Medicine, Isehara, Japan

**Keywords:** Aging, Gas6/Axl axis, Gut commensal bacteria, Pathobiont, Bacterial translocation, mucosal immunity

## Abstract

Immunosenescence refers to the development of weakened and/or dysfunctional immune responses associated with aging. Several commensal bacteria can be pathogenic in immunosuppressed individuals. Although *Klebsiella pneumoniae* is a commensal bacterium that colonizes human mucosal surfaces, the gastrointestinal tract, and the oropharynx, it can cause serious infectious diseases, such as pneumonia, urinary tract infections, and liver abscesses, primarily in elderly patients. However, the reason why *K. pneumoniae* targets elderly population remains unclear. This study aimed to determine how the intestinal immune response of the host to *K. pneumoniae* varies with age. To this end, the study analyzed an *in vivo K. pneumoniae* infection model using aged mice, as well as an *in vitro K. pneumoniae* infection model using a Transwell insert co-culture system comprised of epithelial cells and macrophages. In this study, we demonstrate that growth arrest-specific 6 (Gas6), released by intestinal macrophages that recognize *K. pneumoniae*, inhibits bacterial translocation from the gastrointestinal tract by enhancing tight-junction barriers in the intestinal epithelium. However, in aging mice, Gas6 was hardly secreted under *K. pneumoniae* infection due to decreasing intestinal mucosal macrophages; therefore, *K. pneumoniae* can easily invade the intestinal epithelium and subsequently translocate to the liver. Moreover, the administration of Gas6 recombinant protein to elderly mice prevented the translocation of orally infected *K. pneumoniae* from the gastrointestinal tract and significantly prolonged their survival. From these findings, we conclude that the age-related decrease in Gas6 secretion in the intestinal mucosa is the reason why *K. pneumoniae* can be pathogenic in the elderly, thereby indicating that Gas6 could be effective in protecting the elderly against infectious diseases caused by gut pathogens.

**Author Summary:** Aging causes a weakened/dysfunctional human immune system, reducing the ability to combat pathogens. Understanding the molecular mechanisms underlying age-related immunosenescence is critical for the development of preventive therapies against bacterial infectious diseases in elderly individuals. *Klebsiella pneumoniae* is a representative “pathobiont” that causes serious systemic infections such as pneumonia, urinary tract infections, and liver abscesses, mainly in the elderly. However, it remains unclear how *K. pneumoniae* targets the elderly population. Here, we show that growth arrest-specific 6 (Gas6), released by intestinal macrophages that recognize *K. pneumoniae*, inhibits bacterial invasion into the intestinal epithelium and subsequent translocation to the liver. However, in elderly mice, Gas6 is hardly secreted due to decreased intestinal mucosal macrophages; therefore, *K. pneumoniae* can easily translocate to the liver from the gastrointestinal tract. We concluded that the reason *K. pneumoniae* can be pathogenic to the elderly is the age-related decrease in Gas6 secretion in the intestinal mucosa. Moreover, we revealed that the administration of Gas6 to elderly mice significantly prevented systemic translocation of orally infected *K. pneumoniae*. Our findings provide new insights into the prevention of infectious diseases in the elderly.

## Introduction

Weakened and/or dysfunctional immune responses caused by aging, results in a reduced ability to combat pathogens [1]. Certain infections are more prevalent in the elderly than in younger adults, and there is no doubt that compromised immune function in the elderly is strongly related to a variety of infectious diseases [2]. In particular, the community-acquired pneumonia and urinary tract infections are 3- and 20-fold more common in elderly, respectively [2]. Additionally, the microorganisms that cause infectious disease in the elderly are becoming more diverse; some commensal bacteria that exert pathogenicity in the immunosenescent patients, termed “pathobionts,” can become a fatal clinical concern [3].

*Klebsiella pneumoniae* is an encapsulated gram-negative bacterium found in soil, water, and on the surfaces of medical devices [4]. *K. pneumoniae* is a commensal bacterium that colonizes human mucosal surfaces, the gastrointestinal tract, and the oropharynx. It is a gut pathogen that causes serious infectious diseases such as pneumonia, urinary tract infections, and liver abscesses in immunocompromised patients, including the elderly [4, 5]. In the 1980s, highly virulent strains of *K. pneumoniae* that caused liver infections in healthy individuals were isolated [6, 7]. Additionally, *K. pneumoniae* has become increasingly resistant to antibiotics. In fact, infections caused by carbapenem-resistant *K. pneumoniae* have resulted in a significant increase in morbidity and mortality [8]. Therefore, it is crucial preventing infectious diseases caused by *K. pneumoniae* than developing novel antibiotics. Although the pathogenicity of *K. pneumoniae* needs to be understood in detail to design novel strategies to prevent its infection, the detailed mechanisms of host-*K. pneumoniae* interactions associated with the development of these diseases remain unclear. It is unclear why *K. pneumoniae* specifically targets elderly populations. The aim of this study was to determine how the host’s intestinal immune response to *K. pneumoniae* changes with age. In this context, we analyzed an *in vivo K. pneumoniae* infection model using elderly mice and an *in vitro K. pneumoniae* infection model using a Transwell insert co-culture system based on epithelial cells and macrophages. The results showed that growth arrest-specific 6 (Gas6), released by macrophages that recognize *K. pneumoniae*, enhances the Gas6/Axl axis in the intestinal epithelium. Gas6 is a secreted protein and ligand of the Axl tyrosine kinase receptor. The binding of Gas6 to Axl mediates several biological signals related to cell proliferation, survival, and migration [9]. We showed that Gas6 release in the intestinal mucosa decreases with age, allowing *K. pneumoniae* to readily invade the intestinal epithelium and cause systemic infections. Our findings have the potential to facilitate the development of preventative measures for infectious diseases caused by *K. pneumoniae* in the elderly.

## Results

### *K. pneumoniae* oral infection in elderly mice easily invade the gastrointestinal submucosa and translocate to the liver

To examine age-related differences in susceptibility to *K. pneumoniae*, 15-or 57-week-old mice were given antibiotic-laced drinking water and orally infected with *K. pneumoniae* (**Fig 1A**). To distinguish between endogenous (a gut commensal) *K. pneumoniae* and exogenous (orally infected) *K. pneumoniae*, we electroporated *K. pneumoniae* ATCC43816 with the pmCherry plasmid (**S1A Fig**) and then infected the mice. Infection with *K. pneumoniae* ATCC43816 pmCherry severely reduced the survival of 57-week-old mice compared to 15-week-old mice (**Fig 1B**). At 2 days post-infection, we observed desquamation of the epithelial cells in the cecal mucosal layer (**S1B Fig, black dotted line**) and edema of the cecal submucosa in 57-week-old mice, but not in 15-week-old mice (**S1B Fig, red dotted line**). Moreover, we observed liver damage in 57-week-old mice infected with *K. pneumoniae* ATCC43816 pmCherry but not in 15-week-old mice (**S1B Fig**). The number of bacteria in the cecal mucosa and liver of 57-week-old mice 2 days post-infection was significantly higher than that in 15-week-old mice (**Fig 1C**). Immunostaining with an anti-mCherry antibody detected orally infected *K. pneumoniae* on the surface of cecal epithelial cells in 15-week-old mice; however, these signals in 57-week-old mice were detected within the cecal submucosa, indicating that orally infected *K. pneumoniae* were only able to invade the cecal mucosal layer of elderly mice (**Fig 1D**). Moreover, signals generated by *K. pneumoniae* were much stronger in the livers of 57-week-old mice than in those of 15-week-old mice, indicating that orally infected *K. pneumoniae* in 57-week-old mice translocated easily to the liver (**Fig 1E**).

**Fig 1.**
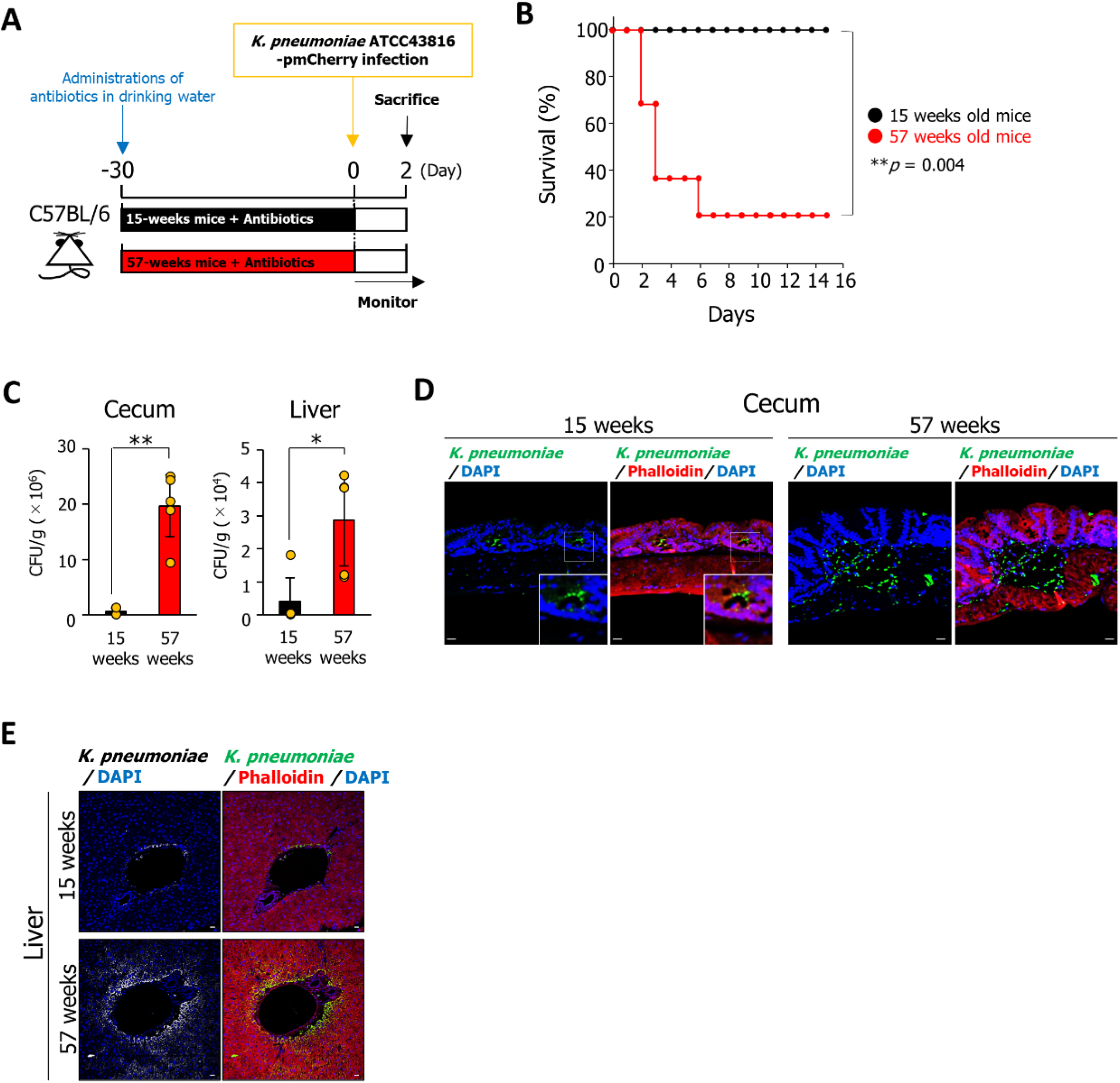
Oral infection with *K. pneumoniae* is highly pathogenic in elderly mice but not in young adult mice. **(A)** Treatment scheme used to analyze age-related differences in susceptibility to *K. pneumoniae*. Mice aged 15-or 57-weeks were administered antibiotics for 4 weeks prior to oral inoculation of *K. pneumoniae* ATCC43816 pmCherry (5 × 10^7^ bacteria). Survival was monitored daily. At 2 days post-infection, the mice were sacrificed, and the cecum and liver were harvested. **(B)** Effect of aging on the survival of mice infected with *K. pneumoniae* ATCC43816 pmCherry. *p*-values were determined using the log-rank test (n = 6 per group). **(C)** Bacterial counts in the cecum and liver were determined 2 days after infection. Cecum and liver tissues were homogenized in PBS, and the homogenates were plated onto LB agar containing 400 μg/mL ampicillin. The number of CFU was determined. Data are presented as the mean ± SD (n = 6 per group). \**p* < 0.05, \*\**p* < 0.01. **(D and E)** Sections of cecal mucosa (D) or liver (E) were from mice aged 15 or 57 weeks at 2 days after infection with *K. pneumoniae* ATCC43816 pmCherry and immunostained with an anti-mCherry antibody and rhodamine phalloidin staining. Scale bar = 20 μm.

### Macrophages upregulate the expression of tight junction proteins between epithelial cells under *K. pneumoniae* infection and prevent bacterial invasion

To determine why *K. pneumoniae* invasion into the intestinal mucosa of young adult mice was inhibited, we constructed an *in vitro K. pneumoniae* infection model using a Transwell insert co-culture system based on Caco-2 epithelial cells and RAW264.7 macrophages (**Fig 2A**). Caco-2 cells co-cultured with RAW264.7 macrophages in the Transwell insert co-culture system were infected with *K. pneumoniae* for 1.5 h. Next, the Caco-2 cells were incubated for 6 h in Dulbecco’s modified Eagle’s medium (DMEM) containing 100 μg/mL gentamycin to kill extracellular bacteria (**Fig 2A**). Hematoxylin and eosin (H&E) staining revealed that Caco-2 cells were not affected by *K. pneumoniae* infection under the co-culture conditions employed; however, in the absence of RAW264.7 macrophages, *K. pneumoniae* ruptured Caco-2 cells (**S2 Fig**). Furthermore, in the absence of RAW264.7, the number of bacteria invading Caco-2 cells was significantly higher than that in the presence of RAW264.7, suggesting that macrophages inhibit bacterial invasion into epithelial cells (**Fig 2B**). Moreover, immunostaining of *K. pneumoniae* reaching the basolateral side of Caco-2 cells was detected in the absence of RAW264.7 macrophages (**Fig 2C**). Expression of tight junction proteins ZO-1 and occludin of Caco-2 cells infected with *K. pneumoniae* in the absence of RAW264.7 macrophages was significantly lower than that in the presence of RAW264.7 macrophages (**Fig 2D and 2E)**.

**Fig 2.**
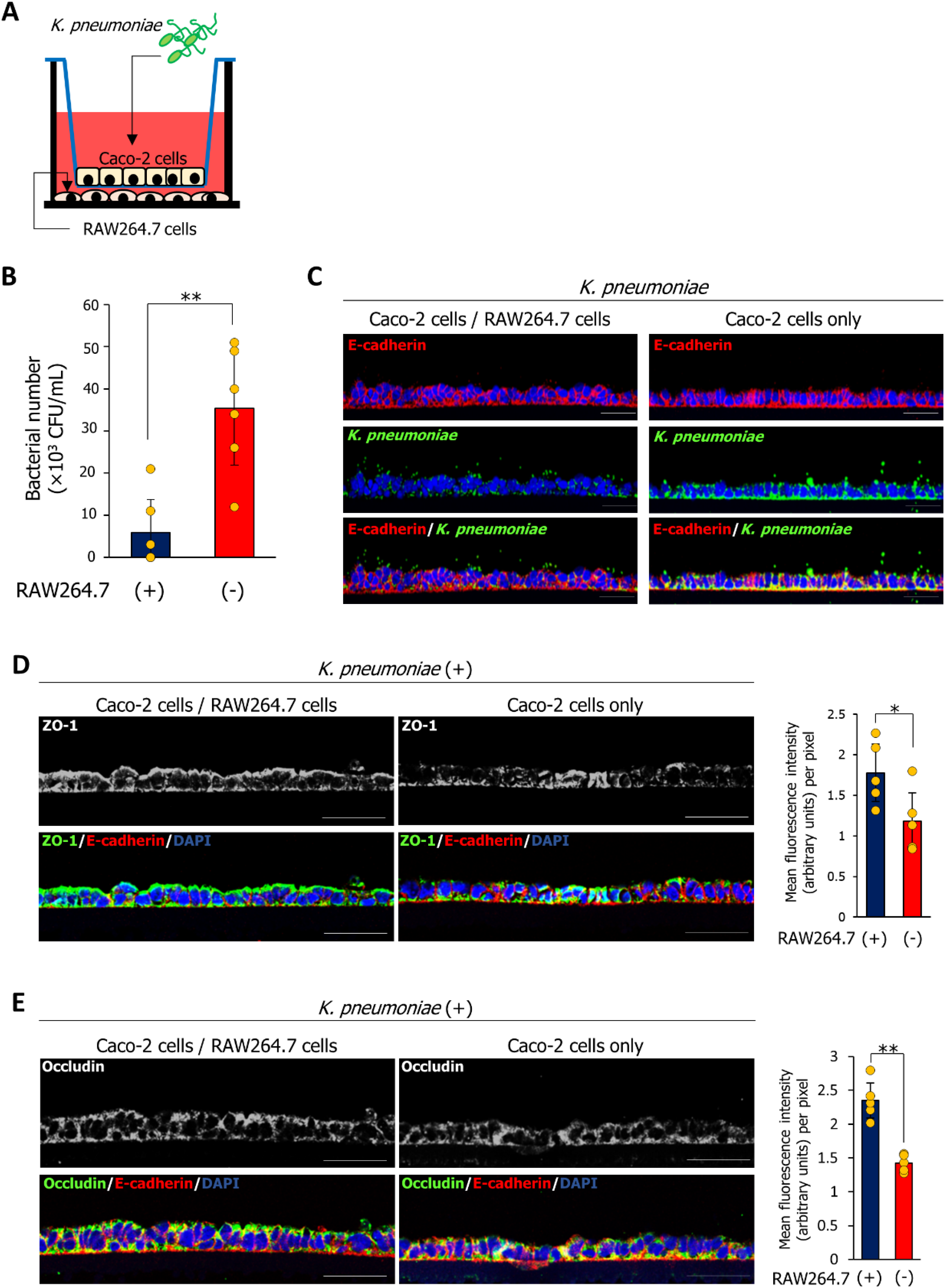
Macrophages inhibit the invasion of intestinal epithelial cells by *K. pneumoniae in vitro*. **(A)** An *in vitro K. pneumoniae* infection model using a Transwell insert co-culture system consisting of Caco-2 cells grown on the insert and RAW264.7 macrophages grown in the cell culture well. **(B)** Bacterial counts in Caco-2 cells. Caco2 cells were lysed with PBS/1% Triton X-100, and the lysate was plated on LB agar. The number of CFU was counted after 24 h incubation. Data are represented as mean ± SD. \*\**p* < 0.01. **(C)** Caco-2 cells grown on the insert in the presence or absence of RAW264.7 macrophages were immunostained with an anti-E-cadherin antibody and an anti-*Klebsiella pneumoniae* antibody. Scale bar = 50 μm. **(D and E)** Caco-2 cells grown on the insert in the presence or absence of RAW264.7 macrophages were immunostained with an anti-E-cadherin antibody, and an anti-ZO-1 antibody (D) or an anti-occludin antibody (E). Staining intensity of ZO-1 (D) or occludin (E) was analyzed by ImageJ software. Data represent the mean ± SD. \**p* < 0.05, \*\**p* < 0.01.

### Gas6 and Axl are released by macrophages that recognize *K. pneumoniae* infection, and the Gas6 is co-localized with the Axl tyrosine kinase receptor on the epithelial cells

To explore how RAW264.7 macrophages regulate the expression of tight junction proteins involved in the repression of bacterial invasion, we focused on cytokines released during *K. pneumoniae* infection. After exposure to *K. pneumoniae* for 1.5 h, Caco-2 cells grown in a Transwell co-culture system with RAW264.7 macrophages were incubated for 6 h with DMEM containing 100 μg/mL gentamycin. The culture medium was harvested for cytokine array analysis (QAM-CAA-4000; RayBiotech, Peachtree Corners, GA, USA). Cytokine array analysis revealed that the *K. pneumoniae* infection significantly increased the release of the receptor tyrosine kinase Axl (**Fig 3A**). To confirm this result, the culture supernatant or the bacterial lysate from *K. pneumoniae* was added to RAW264.7 or Caco-2 cells, and the culture medium was collected for Axl measurement using an ELISA. The secretion of Axl by RAW264.7 macrophages after exposure to *K. pneumoniae* culture supernatant or bacterial lysate increased significantly in a dose-dependent manner (**Fig 3B**). Axl secretion by Caco-2 cells was not observed (**Fig 3B**). Gas6 is an Axl ligand. Gas6/Axl signaling, which is triggered by the binding of Gas6 to Axl, is involved in cell proliferation, survival, and migration [9]. Therefore, we investigated whether *K. pneumoniae* enhanced Gas6 secretion by RAW264.7 macrophages. Gas6 secretion by RAW264.7 macrophages increased significantly in a dose-dependent manner upon exposure to *K. pneumoniae* culture supernatant or bacterial lysate; however, Caco-2 cells did not exhibit an increase in Gas6 secretion (**Fig 3C**). Interestingly, heat-treated (95 °C, 5 min) bacterial supernatants or lysates also induced Gas6 and Axl secretion by RAW264.7, indicating that the Axl- and Gas6-inducing factors produced by *K. pneumoniae* are heat-resistant (**Fig 3B and 3C**). Moreover, we observed co-localization of Gas6 and Axl in Caco-2 cells infected with *K. pneumoniae* in the presence of RAW264.7 macrophages (**Fig 3D**). Co-localization of Gas6 and Axl in Caco-2 cells was not detected in the absence of *K. pneumoniae* or in the presence of *K. pneumoniae* without RAW264.7 macrophages (**Fig 3D**).

**Fig 3.**
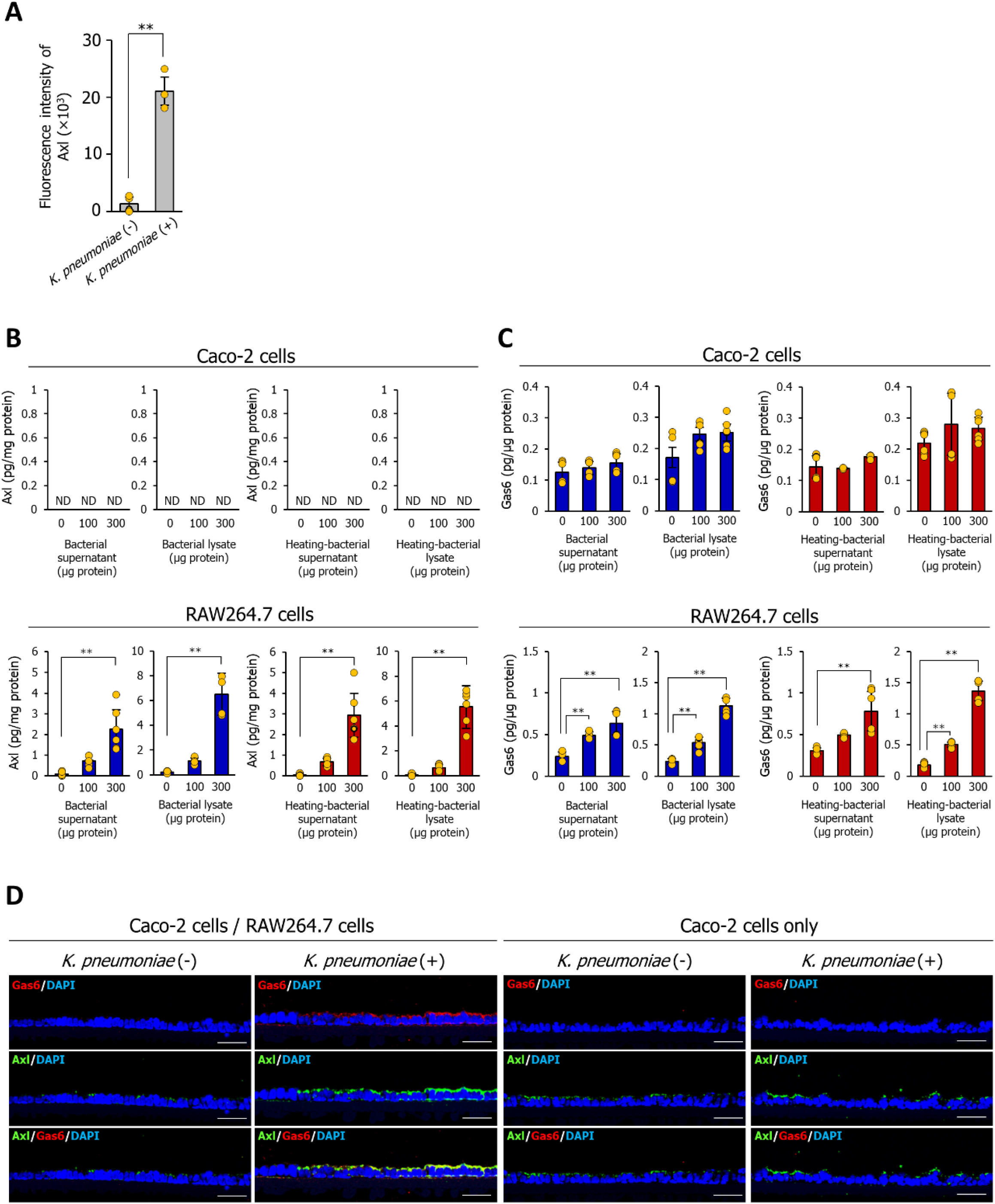
Gas6 released by macrophages that recognized *K. pneumoniae* infection is co-localized with Axl tyrosine kinase receptor on the epithelial cells. **(A)** Cytokine array analysis of the *K. pneumoniae* infection model based on the Transwell insert co-culture system consisting of Caco-2 cells and RAW264.7 macrophages. Quantification of Axl signals using a laser scanner. Data are presented as the mean ± SD. \*\**p* < 0.01. **(B and C)** Secretion of Axl (B) and Gas6 (C) by Caco-2 cells or RAW264.7 macrophages. Culture supernatants or lysates of *K. pneumoniae* were added to Caco-2 cells or RAW264.7 macrophages for 12 h. The culture media were then collected for ELISA analysis of Axl or Gas6 levels. Data are presented as the mean ± SD. \*\**p* < 0.01. **(D)** *K. pneumoniae*-infected Caco-2 cells grown on a Transwell insert in the presence or absence of RAW264.7 macrophages were immunostained with anti-Gas6 and anti-Axl antibodies. Scale bar = 50 μm.

### Gas6 released by macrophages, prevents bacterial invasion by upregulating the expression of tight junction proteins between epithelial cells

Based on these results, we hypothesized that the released Gas6 prevents bacterial invasion by increasing the expression of ZO-1 and occludin in Caco-2 cells via binding to Axl. To test this, Caco-2 cells were treated with an Axl inhibitor (R428; 20 nM) or 1 μg of anti-Gas6 antibody for 3 h prior to infection with *K. pneumoniae*. Upon *K. pneumoniae* infection, the expression of Axl, Gas6, ZO-1, and occludin in Caco-2 cells was increased by the presence of RAW264.7 macrophages (**Fig 4A, lanes 1 and 2**). The increased expression of Axl, ZO-1, and occludin was inhibited by Axl inhibitor (R428) or anti-Gas6 antibody (**Fig 4A, lanes 4 and 6**). Immunostaining analysis of Caco-2 cells infected with *K. pneumoniae* in the presence of RAW264.7 macrophages revealed a significant decrease in the expression of ZO-1 and occludin in the presence of the Axl inhibitor or anti-Gas6 antibody (**Fig 4B**). Next, we examined the effect of an Axl inhibitor or an anti-Gas6 antibody on the invasion of *K. pneumoniae* into Caco-2 cells. The presence of RAW264.7 significantly inhibited the invasion of *K. pneumoniae* within Caco-2 cells, and this effect was abolished upon treatment with an Axl inhibitor or anti-Gas6 antibody (**Fig 4C**). These results suggest that macrophage-released Gas6 enhances the tight junction barrier between epithelial cells to prevent *K. pneumoniae* invasion.

**Fig 4.**
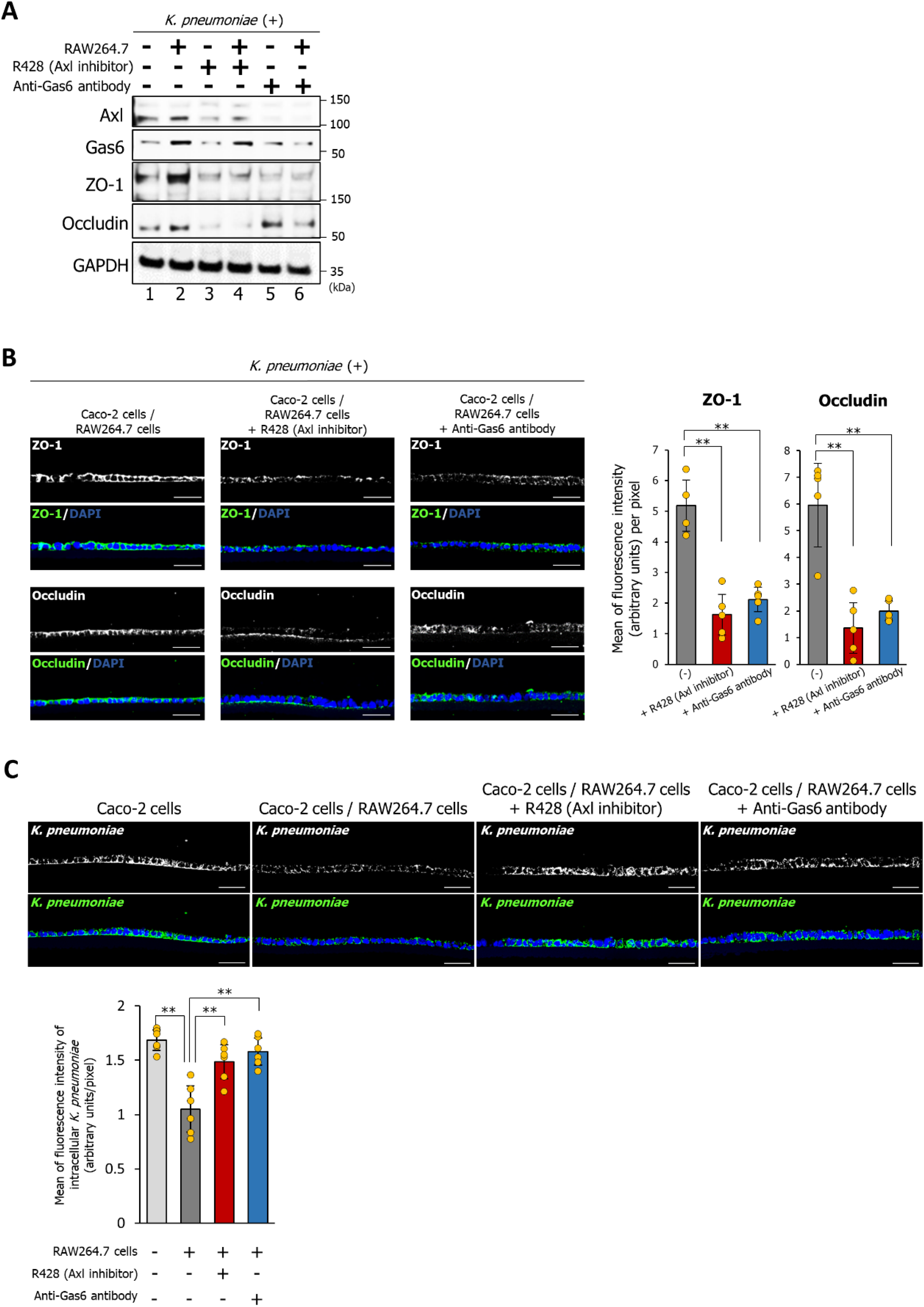
Gas6/Axl signaling in Caco-2 cells inhibits *K. pneumoniae* invasion by enhancing the expression of tight junction proteins. **(A)** Western blotting was performed to detect the expression of Axl, Gas6, ZO-1, and occludin in Caco-2 cells infected with *K. pneumoniae* in the presence of an Axl inhibitor (R428) or an anti-Gas6 antibody. Prior to *K. pneumoniae* infection, cells were treated for 3 h with Axl inhibitor (R428; 20 nM) or 1 μg of anti-Gas6 antibody. **(B and C)** Axl inhibitor (R428; 20 nM) or 1 μg of anti-Gas6 antibody were added to Caco-2 cells grown on the insert in in the presence or absence of RAW264.7 macrophages for 3 h prior to *K. pneumoniae* infection. Next, Caco-2 cells were immunostained with an anti-ZO-1 antibody (B), an anti-occludin antibody (B), and an anti-*Klebsiella pneumoniae* antibody (C). Scale bar = 50 μm. Immunostaining intensities of ZO-1 (B), occludin (B), and *K. pneumoniae* (C) were analyzed by ImageJ software. Data are presented as the mean ± SD. \*\**p* < 0.01.

### Expressions of Gas6 and Axl in the cecal mucosa under *K. pneumoniae* infection decreased with aging

We then investigated whether the expression levels of Gas6 and Axl in *K. pneumoniae* infection differ with age. Immunostaining of Gas6 and Axl was stronger in the cecal mucosa of 15-week-old mice infected with *K. pneumoniae* ATCC43816 pmCherry than that in 57-week-old mice, and Gas6 and Axl co-localized in cecal epithelial cells (**Fig 5A**). Interestingly, co-localization was detected at both the apical and basolateral sides of the epithelial cells (**Fig 5A**). In contrast, Gas6 was barely detectable in the liver epithelial cells of 15-week-old or 57-week-old mice infected with *K. pneumoniae* ATCC43816 pmCherry (**Fig 5B**). Western blotting confirmed a significant decrease in the expression of Axl, Gas6, ZO-1, and occludin in the cecal mucosa of 57-week-old mice compared to that in 15-week-old mice infected with *K. pneumoniae* ATCC43816 pmCherry (**Fig 5C**). However, in the liver, the expression of these molecules did not decrease with age (**Fig 5C**). In addition, we found a significant linear correlation between Gas6 and Axl expression in the cecal mucosa of both 15-week-old and 57-week-old mice (15-week-old mice; *r* = 0.957, 57-week-old mice; *p* = 0.0106, and *r* = 0.996, *p* = 0.0003), but not in the liver (**Fig 5D**). Our findings suggest that age-related decrease in the expression of Gas6 and Axl in the intestinal mucosa increases susceptibility to *K. pneumoniae* oral infection. The number of macrophage precursors and macrophages in the bone marrow of the elderly are reported to be significantly reduced [10, 11]. We then examined how the proportion of intestinal mucosal macrophages changes with age. The proportion of F4/80+ macrophages in the intestinal mucosa of aged mice (56-week-old) was 8.87%, whereas it was 29.5% in young mice (14-week-old) (**Fig 5E**). These results suggest that an age-related decrease in intestinal mucosal macrophages may be one of the reasons for reduced Gas6 expression in elderly mice infected with *K. pneumoniae*.

**Fig 5.**
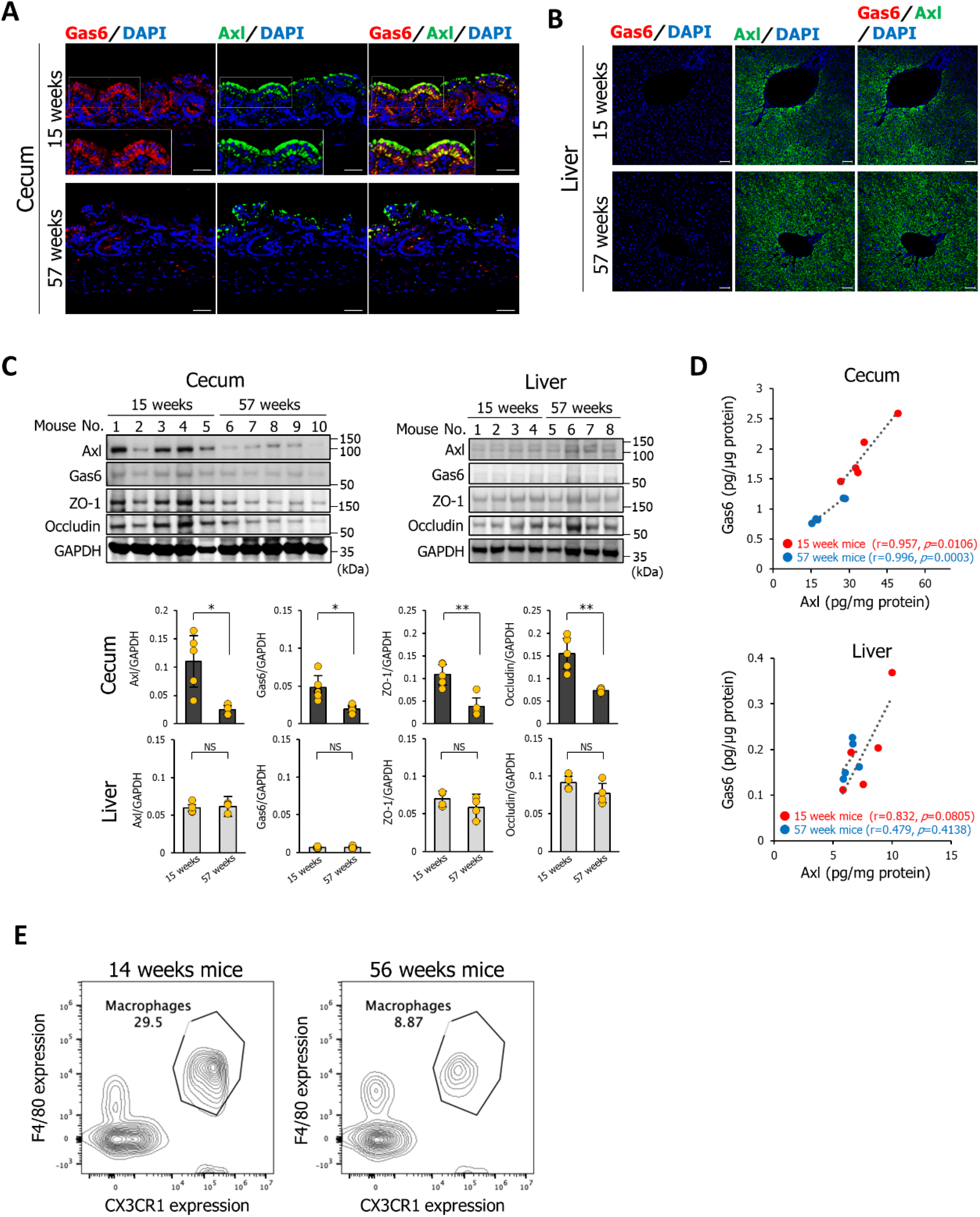
Gas6 and Axl expression in the cecal mucosa during *K. pneumoniae* infection decreases with aging. **(A and B)** Sections of cecal mucosa (A) or liver (B) were from mice aged 15 or 57 weeks at 2 days after infection with *K. pneumoniae* ATCC43816 pmCherry and immunostained with an anti-Gas6 antibody and an anti-Axl antibody. Scale bar = 50 μm. **(C)** Detection of Axl, Gas6, ZO-1, and occludin by western blotting. Cecum and liver tissues were collected, homogenized, and analyzed using an anti-Axl antibody, an anti-Gas6 antibody, an anti-ZO-1 antibody, or an anti-occludin antibody. Signal intensity was analyzed by ImageJ software. Data are presented as the mean ± SD (n = 5 per group). NS: not significant, \**p* < 0.05, \*\**p* < 0.01. **(D)** Linear correlation between Gas6 expression and Axl expression in the cecum or liver of mice aged 15 (red circles) or 57 (blue circles) weeks infected with *K. pneumoniae* ATCC43816 pmCherry. *r* > 0.70 and *p* < 0.05. **(E)** The population of CD11b+F4/80+ macrophages in the intestinal mucosa of young (14-week-old) and old (56-week-old) mice was examined by staining with anti-F4/80 and anti-CD11b antibodies. The percentage of F4/80-positive/CX3CR1-negative cells in 14-week-old and 56-week-old mice was 29.5% and 8.87%, respectively. Data are representative of four mice.

### Administration of Gas6 prevents systemic infection by orally infecting *K. pneumoniae* in elderly mice

Based on our findings, we hypothesized that the administration of Gas6 would inhibit bacterial invasion of epithelial cells. To test this hypothesis, recombinant Gas6 protein was exposed to the apical or basolateral sides of Caco-2 cells grown in a Transwell plate for 18 h (**Fig 6A**). Regardless of the side exposed to Gas6 recombinant protein, the expression of Axl, ZO-1, and occludin increased in Caco-2 cells stimulated with Gas6 (**Fig 6B**). Next, we added Gas6 recombinant protein (1 μg) to the apical side of Caco-2 cells grown in transwell co-culture systems for 3 h prior to *K. pneumoniae* infection. After 1.5 h of infection with *K. pneumoniae*, the cells were incubated for 6 h with DMEM containing 100 μg/mL gentamycin and 1 μg Gas6 recombinant protein to kill extracellular bacteria. Administration of the Gas6 recombinant protein increased the expression of Axl, ZO-1, and occludin in Caco-2 cells in the absence of *K. pneumoniae* (**Fig 6C, lanes 1 and 2**). Increased expression of these proteins induced by the Gas6 recombinant protein was also detected in Caco-2 cells infected with *K. pneumoniae* (**Fig 6C, lanes 3 and 5**). Immunostaining analysis revealed that the expression of ZO-1 and occludin (but not E-cadherin) by *K. pneumoniae*-infected Caco-2 cells in the absence of RAW264.7 macrophages increased significantly after treatment with Gas6 recombinant protein (**Fig 6D**). Furthermore, treatment with Gas6 recombinant protein significantly decreased the signals of *K. pneumoniae* immunostaining within Caco-2 cells (**Fig 6E**). These findings show that administration of Gas6 instead of RAW264.7 macrophages can repress bacterial invasion of intestinal epithelial cells by enhancing tight junction barriers.

**Fig 6.**
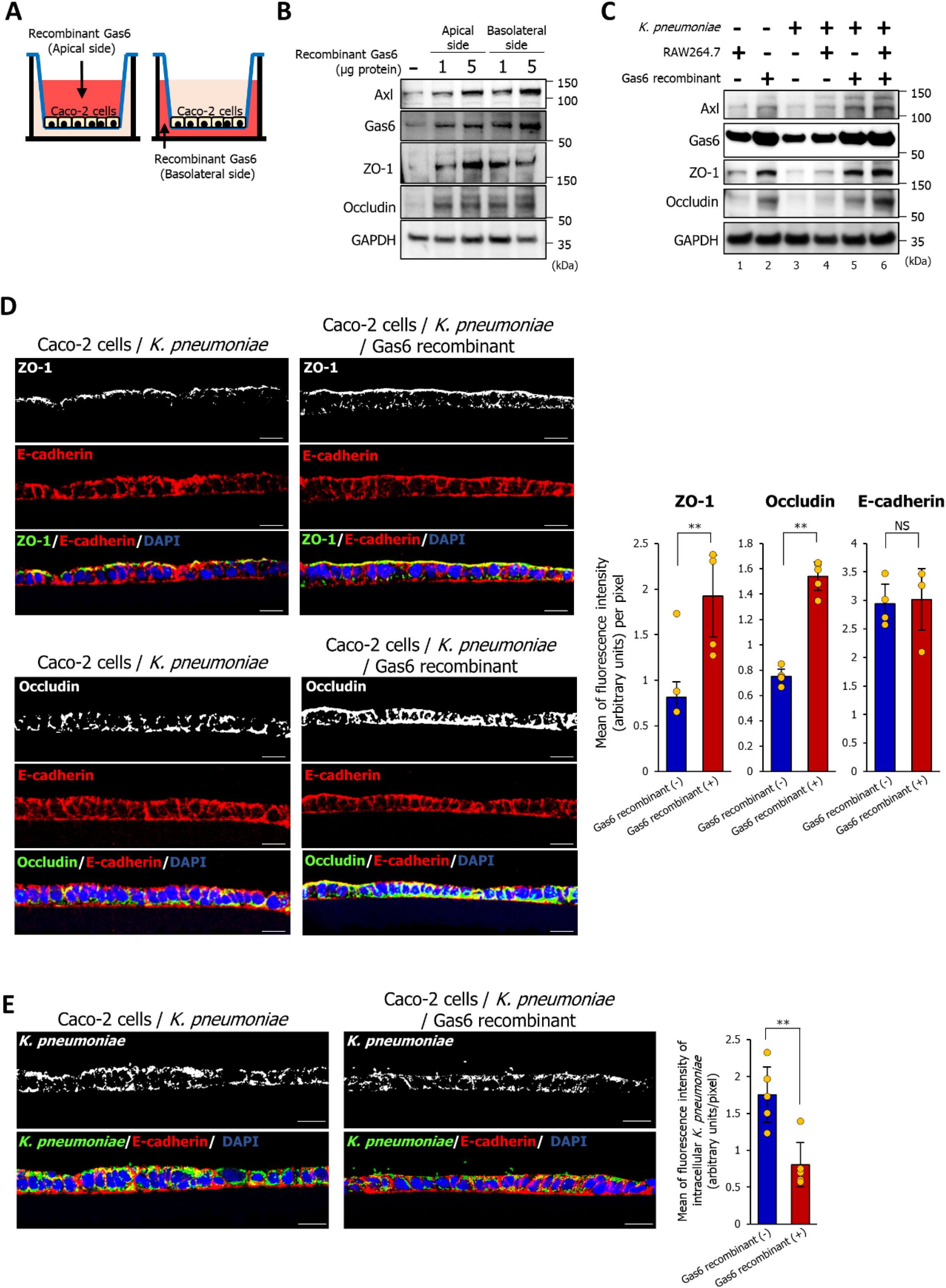
Gas6 represses the invasion of *K. pneumoniae* into Caco-2 cells by increasing the expression of ZO-1 and occludin. **(A)** Administration of Gas6 recombinant protein to Caco-2 cells to analyze the expression of Axl, ZO-1, and occludin. Addition of Gas6 recombinant protein to the apical surface (left scheme) or the basolateral side (right scheme) of Caco-2 cells. **(B)** Western blot analysis to detect Axl, Gas6, ZO-1, and occludin in Caco-2 cells treated with human Gas6 recombinant protein from the apical or basolateral sides. **(C)** Western blot analysis was performed to detect Axl, Gas6, ZO-1, and occludin. Gas6 recombinant protein (1 μg) was added to the apical side of Caco-2 cells grown in a Transwell co-culture system for 3 h prior to *K. pneumoniae* infection. **(D and E)** Gas6 recombinant protein (1 μg) was added to Caco-2 cells grown on the Transwell insert for 3 h prior to *K. pneumoniae* infection. Next, Caco-2 cells were immunostained with an anti-ZO-1 antibody (C), an anti-occludin antibody (C), an anti-*Klebsiella pneumoniae* antibody (D), and an anti-E-cadherin antibody. The signal intensities of ZO-1, occluding, E-cadherin, and *K. pneumoniae* were analyzed by ImageJ software. Data are presented as the mean ± SD. NS: not significant, \*\**p* < 0.01.

Next, we investigated whether Gas6 administration protects elderly mice against *K. pneumoniae* infection. Prior to bacterial infection, 57-week-old mice received three intraperitoneal injections of Gas6 recombinant protein (125 μg protein/kg) every 24 h (**Fig 7A**). There was no significant difference between the survival influences of 15-week-old mice and Gas6-treated 57-week-old mice upon infection with *K. pneumoniae*, indicating that Gas6 recombinant proteins protect the elderly host against *K. pneumoniae* infectious disease (**Fig 7B**). The desquamation area of the cecal epithelium observed in *K. pneumoniae-*infected 57-week-old mice was barely detectable in Gas6-treated 57-week-old mice (**S3A and S3B Fig)**. While the number of bacteria invading the cecal mucosa and translocating to the liver was significantly higher in 57-week-old mice than in 15-week-old mice, the increase was significantly attenuated by the administration of Gas6 recombinant protein (**Fig 7C**). The expression levels of Axl, ZO-1, and occludin in the cecal mucosa of Gas6-treated 57-week-old mice infected with *K. pneumoniae* were restored to levels similar to those in 15-week-old mice (**Fig 7D**). These results revealed the protective potential of Gas6 against *K. pneumoniae* infection in elderly hosts.

**Fig 7.**
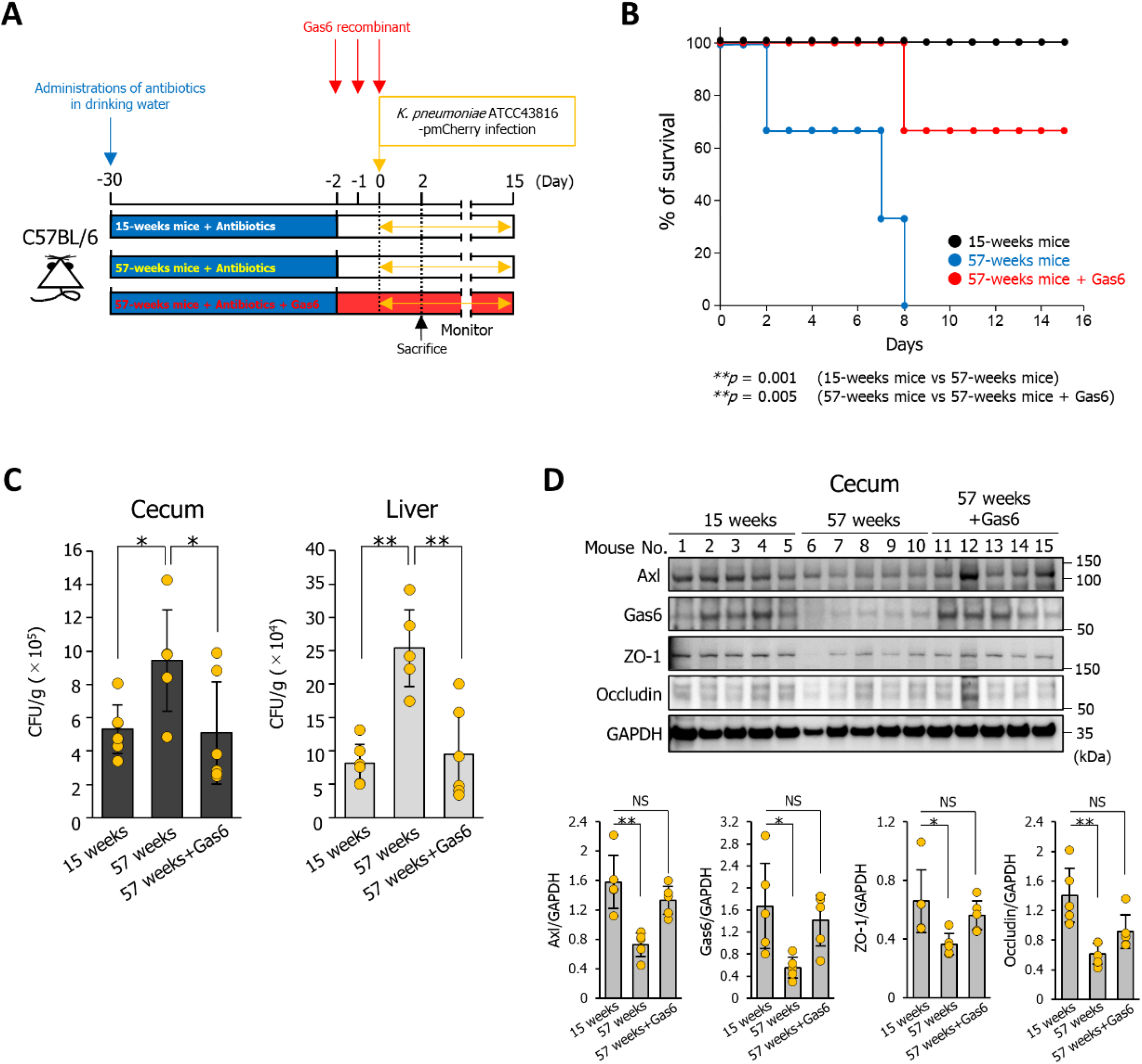
Gas6 prevents systemic infection by orally infecting *K. pneumoniae* in elderly mice. **(A)** Treatment scheme used to analyze the effect of Gas6 recombinant protein on the susceptibility of 57-week-old mice to infection by *K. pneumoniae*. Antibiotics were administered 4 weeks before administration of Gas6 recombinant protein. Gas6 recombinant protein (125 μg protein/kg) was administered intraperitoneally to 57-week-old mice three times every 24 h prior to bacterial infection. **(B)** Effect of Gas6 recombinant protein on survival of 57-week-old mice infected with *K. pneumoniae* ATCC43816 pmCherry. Each mouse was orally inoculated with *K. pneumoniae* ATCC43816 pmCherry (5 × 10^7^ bacteria). *p*-values were determined using the log-rank test (n = 6 per group). **(C)** Bacterial counts in cecum and liver tissues were determined 2 days post-infection. Cecum and liver tissues were homogenized in PBS. The homogenates were plated on LB agar containing 400 μg/mL ampicillin and the number of CFU was counted. Data are presented as the mean ± SD. (n = 6 per group). \**p* < 0.05, \*\**p* < 0.01. **(D)** Detection of Axl, Gas6, ZO-1, and occludin by western blotting. Cecum tissues were collected, homogenized, and analyzed by western blotting with anti-Axl, anti-Gas6, anti-ZO-1, or anti-occludin antibodies. Signal intensity was analyzed by ImageJ software. Data are presented as the mean ± SD (n = 5 per group). NS: not significant, \**p* < 0.05, \*\**p* < 0.01.

## Discussion

*K. pneumoniae* is a potentially pathogenic gut symbiont that can become pathogenic under specific genetic, environmental, or immunocompromised conditions [12, 13]. However, the environmental or immunological conditions that transform *K. pneumoniae* from a gut symbiont to a pathogenic organism remain unclear. Here, we showed that reduced Gas6/Axl signals with aging in the intestinal mucosa induce the invasion of *K. pneumoniae* into the intestinal epithelium, and these bacteria then easily translocate to the liver.

*K. pneumoniae* caused desquamation of the cecal intestinal epithelium of 57-week-old mice and Caco-2 cells (**S1B and S2 Fig**). According to Nakamoto *et al*. [14], *K. pneumoniae* induces epithelial pore formation, which disrupts the intestinal barrier and allows bacterial translocation. Furthermore, the pore-forming ability of *K. pneumoniae* is considered strain-specific [14]. Recently, hypervirulent strains of *K. pneumoniae* were isolated from patients with liver abscesses; these strains exert pathogenic effects even in immune-competent healthy individuals [15]. Conserved genes responsible for pathogenic pore-forming ability have not been identified; therefore, it is unclear whether the hypervirulent strains and the ATCC43816 strain used in this study encode pathogenic genes related to pore-forming capacity. Although further investigations, including the identification of pore-forming toxins, are needed to identify the precise mechanisms by which *K. pneumoniae* induces epithelial injury, the current findings indicate that age-related changes in host-bacterial interactions play a significant role in the induction of *K. pneumoniae*-mediated epithelial cell injury and subsequent bacterial translocation to the liver.

We found a significant linear correlation between the expression levels of Gas6 and Axl in the cecal mucosa (**Fig 5D**). In contrast, this linear correlation and reduced expression of tight junction proteins were not observed in the liver (**Fig 5D**). These results suggest that the mechanism that regulates Gas6/Axl signaling differs between the cecal mucosa and the liver. Liver-resident macrophages (Kupffer cells) are estimated to not secrete Gas6 (or very low amounts); in fact, Gas6 expression in the liver was barely detectable by immunostaining (**Fig 5B and 5C**). Additionally, it is also conceivable that *K. pneumoniae* translocated to the liver may not release a factor that increases Gas6 production by Kupffer cells. Our findings indicate that macrophage-dependent Gas6/Axl signaling in the intestinal epithelium is essential for maintaining the intestinal mucosal barrier.

Previously, *K. pneumoniae* has been known as the “gut pathobiont” that causes serious infections, primarily in immunocompromised individuals. However, multidrug-resistant and hypervirulent strains of *K. pneumoniae* have been identified, which are capable of causing untreatable infections in healthy individuals [16-19]. Thus, a detailed understanding of the biology underlying infectious behavior is necessary to develop new treatment and prevention strategies. The present study shows that the administration of exogenous Gas6 protects elderly mice against *K. pneumoniae* infection by preventing reductions in ZO-1 and occludin expression in the intestinal mucosa (**Fig 7**). Axl is expressed not only on the apical but also on the basolateral cell surface membrane [20]. *In vitro* experiments also showed that the addition of exogenous Gas6 to the apical or basolateral side induced ZO-1 and occludin expression (**Fig 6B**). Taken together, these results indicate that different routes of administration can be considered when using a Gas6 or Gas6 mimic compound to modulate Gas6/Axl signaling in the intestinal epithelium. Our findings provide new insights into strategies to combat *K. pneumoniae* infectious diseases in the elderly.

## Methods

### Ethics statement

All animal experiments were approved by the Keio University (Tokyo, Japan) Animal Research Committee (no. 19048) and the Tokai University (Kanagawa, Japan) Animal Research Committee (no. 222002) and were conducted in accordance with the “Act on Welfare and Management of Animals of Japan,” “Standards relating to the Care and Keeping and Reducing Pain of Laboratory Animals,” “Standards relating to the Methods of Destruction of Animals,” “Guidelines for Proper Conduct of Animal Experiments,” and “Fundamental Guidelines for Proper Conduct of Animal Experiments.”

### Reagents and antibodies

An Axl inhibitor (R428) (Abcam, Cambridge, UK, cat# ab141364), recombinant human Gas6 protein (R&D Systems, Minneapolis, MN, USA, cat# 885-GSB), recombinant mouse Gas6 protein (R&D Systems, cat# 986-GS) (used in administration experiments in mice), and an anti-Gas6 antibody (R&D Systems, cat# AF986) were used as Gas6/Axl signal modulators. Rhodamine phalloidin (Invitrogen, Waltham, MA, USA, R415) was used to stain the polymerized actin. The following antibodies were used for immunostaining: anti-*Klebsiella pneumoniae* (Thermo Fisher Scientific, Cleveland, OH, USA, cat# PA1-7226), anti-E-cadherin (BD Biosciences, San Jose, CA, USA, cat# 610181), anti-ZO-1 (Invitrogen, cat# 61-7300), anti-occludin (Invitrogen, cat# 711500), anti-Axl (GeneTex Inc., Irvine, CA, USA, cat# GTX129407), anti-Gas6 (R&D Systems, cat# AF986), and anti-mCherry (Invitrogen, cat# M11217). The following antibodies were used for western blotting: anti-Axl (GeneTex Inc., cat# GTX129407), anti-Gas6 (R&D Systems, cat# AF986), anti-ZO-1 (Invitrogen, cat# 61-7300), anti-occludin (Invitrogen, cat# 711500), and anti-GAPDH (Cell Signaling Technologies, Danvers, MA, USA, cat# 2118S). The following antibodies were used for flow cytometry analysis: FITC-conjugated anti-CD19 (6D5) (BioLegend, San Diego, CA, USA, cat# 115505), APC-conjugated anti-CD11b (M1/70) (BioLegend, cat# 101211), PerCP-Cy5.5-conjugated anti-CD3 (17A2) (BioLegend, cat# 100217), and PE-Cy7-conjugated anti-F4/80 (BM8) (BioLegend, cat# 123113).

### Cell and bacterial culture

Caco-2 cells, which were purchased from the European Collection of Cell Cultures (ECACC 86010202), were cultured in DMEM (Gibco, Waltham, MA, USA, cat# 11965092) supplemented with 10% fetal bovine serum (FBS), 1% non-essential amino acids (Gibco, cat# 11140-050), and 2 mM L-glutamine (Gibco, cat# 25030-081). RAW264.7 cells purchased from the American Type Culture Collection (ATCC TIB-71) were maintained in DMEM supplemented with 10% FBS. *K. pneumoniae* ATCC43816 was cultured overnight at 37 °C on Luria–Bertani (LB) agar (Nacalai Tesque Inc., Kyoto, Japan, cat# 20067-85). Bacterial counts were determined by measuring the optical density of bacterial suspensions at 550 nm.

### Mice

All the male mice were bred and maintained under specific pathogen-free (SPF) conditions. Six-week-old C57BL/6J male mice were purchased from SLC Japan, Inc. (Shizuoka, Japan) or CLEA Japan, Inc. (Osaka, Japan) and used in experiments at 14 or 15 weeks of age. C57BL/6J mice aged 48–55 weeks were purchased from SLC Japan or CLEA Japan and were used at 57 weeks of age. A previous study reported that the administration of four antibiotics (ampicillin, metronidazole, neomycin, and vancomycin) to mice excluded endogenous short-chain fatty acids (SCFAs) by completely eradicating gut commensal bacteria [21]. To exclude the effects of the gut microbiota, 11 or 53-week-old mice were provided with drinking water containing 1 g/L ampicillin (Sigma-Aldrich, St. Louis, MO, USA, cat# A0166), 1 g/L metronidazole (Sigma-Aldrich, cat# M1547), 1 g/L neomycin (Sigma-Aldrich, cat# N1876), and 0.5 g/L vancomycin (Wako, Osaka, Japan, cat# 222-01303) for 4 weeks [21-23]. Mice that lost more than 20% of their body weight because of their refusal to drink the antibiotic cocktail were excluded from the experiments in accordance with the guidelines of the Keio University Animal Research Committee (no. 19048) or the Tokai University Animal Research Committee (no. 222002).

### Isolation of murine macrophages and Flow cytometry

Intestinal lamina propria cells were isolated from 14 or 56-week-old mice, as described previously with slight changes [24]. The harvested intestine was cut into 10-mm long segments, opened longitudinally, and washed in phosphate-buffered saline (PBS). The washed segments were treated for 10 min at room temperature with 30 mM EDTA. These intestinal pieces were washed thoroughly with PBS and digested for 30 min at 37 °C with 0.5 mg/mL collagenase D (Merck/Sigma-Aldrich, cat#11088858001) in RPMI medium (Gibco, cat#11875093) supplemented with 2% fetal calf serum (FCS). After repeated pipetting, the supernatant was collected after 2 min and passed through a cell strainer (70 μm). The remaining tissue fragments were subjected to a second collagenase digestion, and these steps were repeated until no gross tissue fragments were visible. The digested tissues were washed with 10 mL of PBS, resuspended in 5 mL of 40% Percoll (GE Healthcare, Chicago, IL, USA), and underlaid with 5 mL of 80% Percoll (GE Healthcare) in a 15 mL tube. The Percoll gradient separation was performed by centrifugation at 800 × *g* for 15 min at room temperature. Lymphocytes were collected from the interface of the Percoll gradient and washed with RPMI1640 containing 10% FBS. For cell surface staining, 2 × 106 cells were stained with fluorescence-conjugated anti-F4/80, anti-CD11b, anti-CD19, and anti-CD3e antibodies for 35 min on ice in the presence of an anti-CD16/32 (2.4G2) antibody for blocking. The cells were analyzed using a Cytoflex flow cytometer (Beckman Coulter, Tokyo, Japan). The collected data were analyzed using Flowjo software (Tree Star, Ashland, Oregon, USA). Antibody dilutions were 200-fold for surface staining.

### *K. pneumoniae* infection of mice

To distinguish between intestinal commensal *K. pneumoniae* strains and orally infected *K. pneumoniae* ATCC43816 strains, a pmCherry plasmid (Clontech, Palo Alto, CA, USA, cat #632522) was electroporated into *K. pneumoniae* ATCC43816 using a MicroPulser (Bio-Rad, Hercules, CA, USA). SPF mice aged 11 or 53 weeks were provided with drinking water containing four antibiotics (ampicillin, metronidazole, neomycin, and vancomycin) for 4 weeks prior to *K. pneumoniae* infection. Administration of these antibiotics was stopped prior to bacterial infection. Mice were orally inoculated with *K. pneumoniae* (5 × 10^7^ bacteria) and their survival was monitored daily. To assess the number of bacteria in cecum and liver, *K. pneumoniae*-infected mice were sacrificed 2 days’ post-infection, and tissues were harvested, suspended in PBS, and homogenized with a Qiagen TissueLyser (Qiagen, Hilden, Germany). Serial dilutions of the homogenates were plated on LB agar containing 50 μg/mL ampicillin, and colony-forming units (CFUs) were counted after 24 h of incubation.

Sections (4 μm) of the cecum or liver collected from mice infected with *K. pneumoniae* (5 × 10^7^ bacteria) for 2 days were fixed overnight in 10% formalin neutral buffer solution and embedded in paraffin. The sections were then deparaffinized, and stained with H&E. Desquamation of the epithelial cells or edema of the cecal submucosa was examined using an OPTIKA B-290TB Digital Microscope (Optika Microscope, Milano, Italy). The length of desquamated cecal epithelial cells on H&E-stained specimens was measured using the ImageJ analysis software (National Institute of Health). For immunohistochemistry analysis, tissue sections were treated as described above and incubated overnight at 4 °C with antibodies specific for mCherry (Invitrogen), Gas6 (R&D Systems), or Axl (GeneTex Inc.), followed by incubation for 1 h with DAPI and Alexa Fluor 488-conjugated anti-rat, anti-rabbit, or Alexa Fluor 568-conjugated anti-mouse IgG secondary antibodies. Fluorescence images were obtained using a LSM700 confocal microscope (Carl Zeiss, Oberkochen, Germany). For western blotting, cecum or liver samples collected from each mouse were homogenized in RIPA buffer containing protease inhibitors. Total protein (10 μg/lane) was separated on 10% Bis-Tris Plus gels (Thermo Fisher Scientific, NW00107BOX) and transferred to polyvinylidene difluoride membranes (Amersham, Braunschweig, Germany, cat# 10600122). An anti-GAPDH antibody (Cell Signaling Technologies) was used as a loading control. The intensity of each band was measured using ImageJ software (National Institute of Health, Bethesda, MD, USA).

### *In vitro K. pneumoniae* infection model of Caco-2 cells grown in Transwell culture systems

Caco-2 cells (1 × 10^6^ cells/well) were seeded onto 6-Transwell insert culture plates (Corning, Lowell, MA, USA, cat# 3412). RAW264.7 macrophages were seeded onto 6-well flat-bottom cell culture plates at 1 × 10^5^ cells per well. To establish a co-culture system for Caco-2 cells and RAW264.7 macrophages, inserts on which Caco-2 cells were grown were placed in the wells of culture plates containing RAW264.7 macrophages. RAW264.7 macrophages were primed for 6 h with 1 μg/mL LPS prior to *K. pneumoniae* infection. The apical side of Caco-2 cells grown on Transwell inserts were exposed for 3 h to 20 nM Axl inhibitor (R428) (Abcam), 1 μg of anti-Gas6 antibody (R&D Systems), or 1 μg of human recombinant Gas6 protein (R&D Systems) prior to *K. pneumoniae* infection. *K. pneumoniae* was resuspended in DMEM and incubated for 1.5 h with Caco-2 cells at a multiplicity of infection of 50. The cells were then washed three times with PBS and incubated for 6 h in DMEM containing 400 μg/mL gentamycin. To measure the number of surviving intracellular cells, cells were washed with PBS and lysed with PBS/1% Triton X-100, followed by plating on LB agar. CFUs were counted after 24 h of incubation. For immunopathological analysis, the transwell membrane was fixed overnight in 4% paraformaldehyde and embedded in paraffin. Sections (4 μm) were deparaffinized, stained with H&E, and incubated overnight at 4 °C with antibodies specific for E-cadherin (BD Biosciences, Franklin Lakes, NJ, USA), *K. pneumoniae* (Thermo Fisher Scientific), ZO-1 (Invitrogen), occludin (Invitrogen), Gas6 (R&D Systems), or Axl (GeneTex Inc.). The sections were then incubated for 1 h with Alexa Fluor 488-conjugated anti-rabbit or Alexa Fluor 568-conjugated anti-mouse IgG. Fluorescence images were obtained using an LSM710 or 700 confocal microscope (Carl Zeiss, Oberkochen, Germany). Quantification of *K. pneumoniae*, E-cadherin, ZO-1, and occludin staining was performed using the ImageJ analysis software (National Institutes of Health). For western blotting, cells were collected from the Transwell membrane using Cell Scraper and treated with RIPA buffer containing protease inhibitors. Total protein was separated (10 μg protein/lane) on 10% Bis-Tris Plus gels (Thermo Fisher Scientific). Caco-2 cells grown on transwell inserts were incubated with or without *K. pneumoniae* for 1.5 h and then incubated for 6 h in DMEM containing 400 μg/mL gentamycin. The culture medium was collected for cytokine array analysis (QAM-CAA-4000; RayBiotech, Peachtree Corners, GA, USA).

### Measurement of Axl and Gas6

Caco-2 and RAW264.7 macrophages (1 × 10^6^ cells per well) were cultured in 6-Transwell insert culture plates (Corning) or 6-well flat-bottom cell culture plates, respectively. *K. pneumoniae* was cultured overnight in DMEM at 37 °C with agitation. After centrifugation (8,000 × *g*, 30 min at 4 °C), the supernatant and bacterial pellets were collected. The bacterial pellets were lysed by sonication. The supernatant and the bacterial lysate were heated at 95 °C for 5 min. Caco-2 cells or RAW264.7 macrophages were incubated overnight at 37 °C with the bacterial supernatants, or lysate. The supernatant was collected, and Axl and Gas6 were detected using human Axl (Abcam, cat# 99976), human Gas6 (Invitrogen, cat# BMS2291), mouse Axl (Abcam, cat# ab100669), or mouse Gas6 (Abcam, cat# ab278087) ELISA kits.

### Statistical analysis

The data are presented as mean ± standard deviation (SD). The means of multiple groups were compared by analysis of variance, followed by Tukey’s tests, using the JSTAT statistical software (version 8.2). Cumulative survival rates were analyzed using the Kaplan–Meier method, and differences in survival between subgroups were assessed using the log-rank test in SPSS version 22 for Windows (SPSS Inc., Chicago, IL, USA). The correlation coefficients (r) and their significance (p) were calculated between the variables using JSTAT statistical software (version 8.2). The animals were randomly assigned to various groups. For repeated immunostaining experiments, the conditions were randomized to account for potential ordering effects. Statistical significance was set at *p* < 0.05.

## Acknowledgments

We are grateful to the Support Center for Medical Research and Education at Tokai University School of Medicine for their technical assistance. This study was supported by the Kao Research Council for the Study of Healthcare Science (C93102) (to H.T.). The infection models in this work were partly supported by a 2021 Tokai University School of Medicine Research Aid grant (to H.T.), Ohyama Health Foundation Inc. (to H.T.), and the Japan Agency for Medical Research and Development (21ck0106701h0001) (to J.M.).

## Author contributions

H.T. conceived the study concept and design, and planned the experiments. H.T., T.O., and J.M. wrote the manuscript. H.T., T.O., R.T., S.T., and J.M. performed experiments. H.T., J.M., H.S., and K.H. analyzed the data.

## Declaration of interests

The authors have no potential conflicts of interest to disclose.

## Notes

### Competing Interest Statement

The authors have declared no competing interest.

